# Distribution and environmental drivers of fungal denitrifiers in global soils

**DOI:** 10.1101/2022.12.06.519296

**Authors:** Yvonne Bösch, Grace Pold, Aurélien Saghaï, Magnus Karlsson, Christopher M. Jones, Sara Hallin

## Abstract

The microbial process denitrification is the primary source of the greenhouse gas nitrous oxide (N_2_O) from terrestrial ecosystems. Fungal denitrifiers, unlike many bacteria, lack the N_2_O reductase and are potential sources of N_2_O. Still, their diversity, distribution, and environmental determinants in terrestrial ecosystems remain unresolved. We used a phylogenetically informed approach to screen 1 980 soil and rhizosphere metagenomes representing 608 globally distributed sampling sites for the denitrification marker gene *nirK*, coding for nitrite reductase. We show that fungal denitrifiers are sparse, yet cosmopolitan and dominated by saprotrophs and opportunistic plant pathogens. Few showed biome-specific distribution patterns. However, members of the *Fusarium oxysporum* species complex, known to produce substantial amounts of N_2_O, were proportionally more abundant and diverse in the rhizosphere than in other biomes. Fungal denitrifiers were most frequently detected in croplands but were most abundant in forest soils. The overall low abundance of fungal relative to bacterial and archaeal denitrifiers suggests that their role in denitrification and contribution to soil N_2_O emissions may be less important than previously suggested. Nevertheless, in relative terms, they could play a role in soils characterized by high carbon to nitrogen ratio and low pH, especially in tundra and boreal and temperate coniferous forests. Our results further indicate that plant-pathogen interactions may favor fungal denitrifiers. Thus, increasing global warming with predicted proliferation of pathogens and the fact that many of the fungi with *nirK* detected in the metagenomes are stress-tolerant cosmopolitans suggest that fungal denitrifier abundance may increase in terrestrial ecosystems.

## INTRODUCTION

Terrestrial ecosystems are major sources of the long-lived stratospheric ozone-depleting substance and greenhouse gas nitrous oxide (N_2_O). Direct emissions from natural soils contribute 58% of the total natural fluxes of N_2_O to the atmosphere (total 9.7 Tg N year^-1^), while agricultural soils account for 32% of the anthropogenic sources (total 7.3 Tg N year^-1^), with global emissions currently increasing by 2% per decade [1]. Nitrous oxide originates primarily from the microbial process denitrification [2, 3]. This process is an alternative to aerobic respiration when oxygen levels are low and reduces nitrate to di-nitrogen (N_2_) in four consecutive reactions. It has predominantly been studied in bacteria, but some archaea and fungi are also known as denitrifiers [4]. Denitrifying fungi are of particular interest, as no species to date has been reported to encode the N_2_O reductase catalyzing the reduction of N_2_O to N_2._ Therefore, fungi terminate denitrification with N_2_O and are potentially important sources of N_2_O from soil. Most fungi for which N_2_O production has been demonstrated belong to the fungal classes Eurotiomycetes and Sodariomycetes, including the genera *Fusarium, Aspergillus, Bionectria*, and *Trichoderma* of which many are putative pathogens [5, 6]. Yet, our limited knowledge about their ecology and distribution in terrestrial biomes, which underpins their capacity to contribute to N_2_O emissions, limits our understanding of their role in terrestrial nitrogen (N) feedbacks to the climate system.

Fungal denitrification has been reported in a range of different terrestrial ecosystems [7–12] although its contribution to total denitrification and N_2_O production varies across these systems [13]. In a few cases, fungal denitrification has been suggested to be more prevalent than bacterial denitrification, e.g. in grasslands, and may increase in agricultural soils depending on management practices [7, 8, 14]. Although increased carbon source complexity [15], lower soil pH [13, 16–19], and low oxygen levels have been shown to favor fungal denitrification [18, 20], broadly conserved edaphic factors selecting for fungal or prokaryotic denitrifiers, if any, are not yet established since current knowledge is based on a limited number of case studies. Therefore, the extant diversity of fungal denitrifiers as well as in which terrestrial habitats fungal denitrifiers thrive, remains uncertain.

Here, we present a comprehensive phylogenetically informed analysis of terrestrial fungal denitrifiers on a global scale by examining their abundance and the distribution of fungal denitrifier genotypes in 1 980 soil and rhizosphere metagenomes representing the major terrestrial biomes (Table S1). Utilizing the *nirK* gene, which encodes the copper-dependent nitrite reductase involved in denitrification, as a marker gene for fungal denitrifiers, we circumvent the current debate about the possible involvement of the fungal nitric oxide reductase P450*nor* in secondary metabolism [21]. The abundance of fungal *nirK* was assessed both per total number of reads and in relation to the overall fungal community based on the fungal 18S rRNA gene counts to determine biome-specific differences as well as edaphic drivers of fungal *nirK* counts and the proportion of fungal denitrifiers within the overall fungal community. We further compared differences across biomes and evaluated the edaphic drivers of the fungal relative to bacterial and archaeal denitrifiers as a measure of their relative capacity for denitrification. The compilation and use of a large, globally distributed dataset of metagenomes and associated metadata allowed us to address the ecology and distribution of fungal denitrifiers across broad environmental gradients and free from PCR-introduced biases. In addition, the updated *nirK* reference phylogeny, including them most comprehensive fungal *nirK* reference to date, used to recruit *nirK* genes from metagenomes is a valuable tool for the community researching the ecology and evolution of denitrification.

## MATERIAL AND METHODS

### Metagenome selection and biome assignment

We searched the literature, National Center for Biotechnology Information (NCBI), and Integrated Microbial Genomes and Microbiomes (IMG/M hosted by the Joint Genome Institute) to construct a database of publicly available soil metagenomes fulfilling the following criteria: (i) sequencing done using Illumina short-read technology, (ii) a minimum of 100 000 reads of at least 150 nucleotides (nt), and (iii) availability of metadata beyond geographic coordinates. The final database consisted of 1 980 metagenomes representing 608 sampling locations across the globe (Fig. 1a; Table S1).

**Fig. 1.**
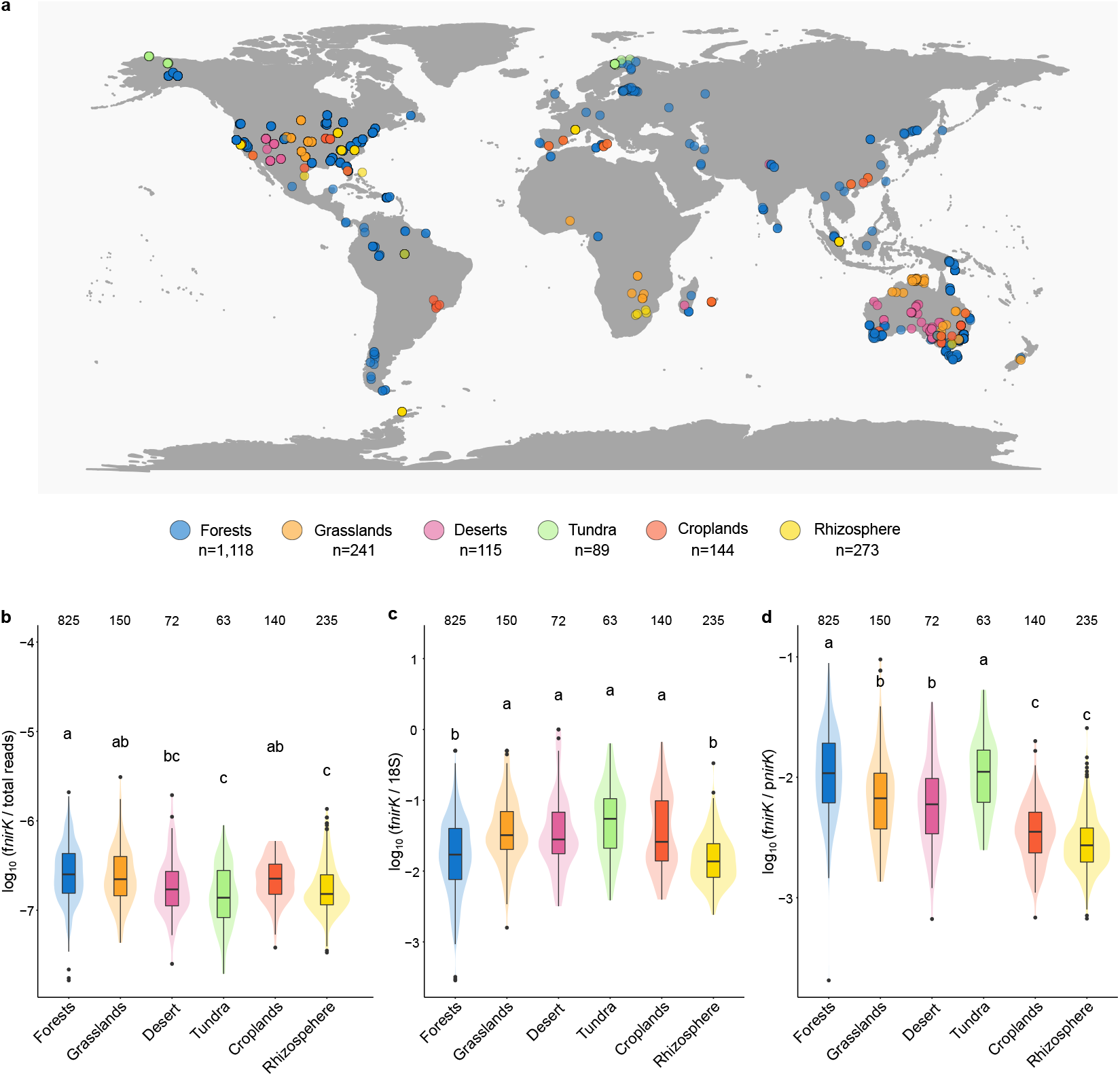
Origin of metagenomes and abundance of fungal *nirK* in terrestrial biomes. (**a**) 1 980 metagenomes representing 608 sampling locations across the globe. The sampling location of 41 rhizosphere samples are not indicated due to the absence of associated geographic coordinates. (**b**) Fungal *nirK* (f*nirK*) gene fragment counts normalized to the total number of reads per metagenome. (**c)** Abundance of f*nirK* relative to fungal 18S rRNA gene (18S) fragment counts. (**d**) Abundance of fungal relative to prokaryotic *nirK* (p*nirK*) gene fragment counts. Different letters indicate significant differences between biomes (ANOVA, Šidák-corrected pairwise comparisons, p <0.05). Numbers above the boxplots in panels **b**-**d** indicate the number of metagenomes within each biome with at least one fungal *nirK* hit. Box limits represent the inter-quartile range (IQR) with median values represented by the centreline. Whiskers represent values ≤1.5 times the upper and lower quartiles, while points indicate values outside this range. The shaded areas show kernel density estimations indicating the distribution of the data.

The metagenomes were classified into three biome levels of increasing ecological complexity using the environment ontology [22] and to discriminate between cropland and non-cropland soils, terrestrial biomes defined by Olson et al., [23] was used. The final collection consisted of 1 980 metagenomes derived from tundra, forests, grasslands and savannas, deserts, croplands, and the rhizosphere of 14 plant taxa (Table S1). Biome assignment was based on the GPS coordinates of each metagenome and performed using the ‘sp’ [24], ‘rgeos’ [25], and ‘rgdal’ [26] packages in R (4.1.2). Level 1 biomes were categorized as terrestrial and host-associated, whilst at Level 2, biomes were distinguished into six terrestrial biomes, which were further subdivided into 13 Level 3 biomes (Table S1). For the host-associated, we restricted our search to plants. Two of the Level 3 biomes (Montane grasslands and shrublands, Tropical and Subtropical coniferous forests) were represented by just two and three samples, respectively, and were excluded from further analysis.

### Generation of *nirK* and 18S rRNA gene phylogenies

We generated a *nirK* reference alignment and phylogeny to identify fungal, bacterial, and archaeal *nirK* sequences within the metagenomes. We first updated and manually curated the alignment of the *nirK* sequences described in Graf et al. [28], consisting of 3 450 sequences extracted from RefSeq genomes (NCBI) available on September 26, 2019. This alignment was converted into a hidden Markov model (HMM) and used to search genome assemblies from NCBI GenBank using *hmmsearch* within the HMMER (v3.2.1) software [29]. Bacterial and archaeal assemblies for this search were downloaded on October 7, 2021, and fungal assemblies on November 21, 2021. Candidate *nirK* amino acid sequences were dereplicated at 100% identity using CD-HIT (v.4.8.1) [30] and aligned against the original seed HMM. The alignments were evaluated and manually refined using a combination of ARB (v.6.1) [31] and FastTree (v.2.1.11) [32] to remove homologous sequences lacking the conserved copper-binding motifs characteristic of the NirK protein [33] and identify an appropriate multicopper oxidase outgroup detected by the HMM search. We also removed sequences originating from genomes that BUSCO (v5.3.1) [34] determined to be > 5% contaminated and/or < 90% complete, except for five > 80% complete Omnitrophica metagenome-assembled genomes (MAGs) for which only medium quality assemblies were available. These bacterial sequences formed a sister clade to the fungi and were thus critical to discriminating fungal versus non-fungal hits. Additional sequences were removed where the taxonomy entered by contributors in NCBI was in a different phylum than that determined by NCBI, or where MMseqs2 taxonomy (v.fcf52600801a73e95fd74068e1bb1afb437d719d) [35] versus the UniRef50 database [36] indicated that *nirK*-containing contigs had inter-domain contamination. Finally, we manually pruned the tree to remove tips with very short terminal branch lengths that would increase the computation time while not allowing better discrimination between fungal and non-fungal gene fragment sequences. The final *nirK* tree was generated with FastTree using an amino acid alignment filter to exclude positions found in fewer than 5% of sequences and the poorly aligned terminal regions. It contains 6 732 sequences (including 373 outgroup sequences) and is comprised of 316 unique fungal *nirK* sequences representing five Ascomycota classes (Fig. 2).

**Fig. 2.**
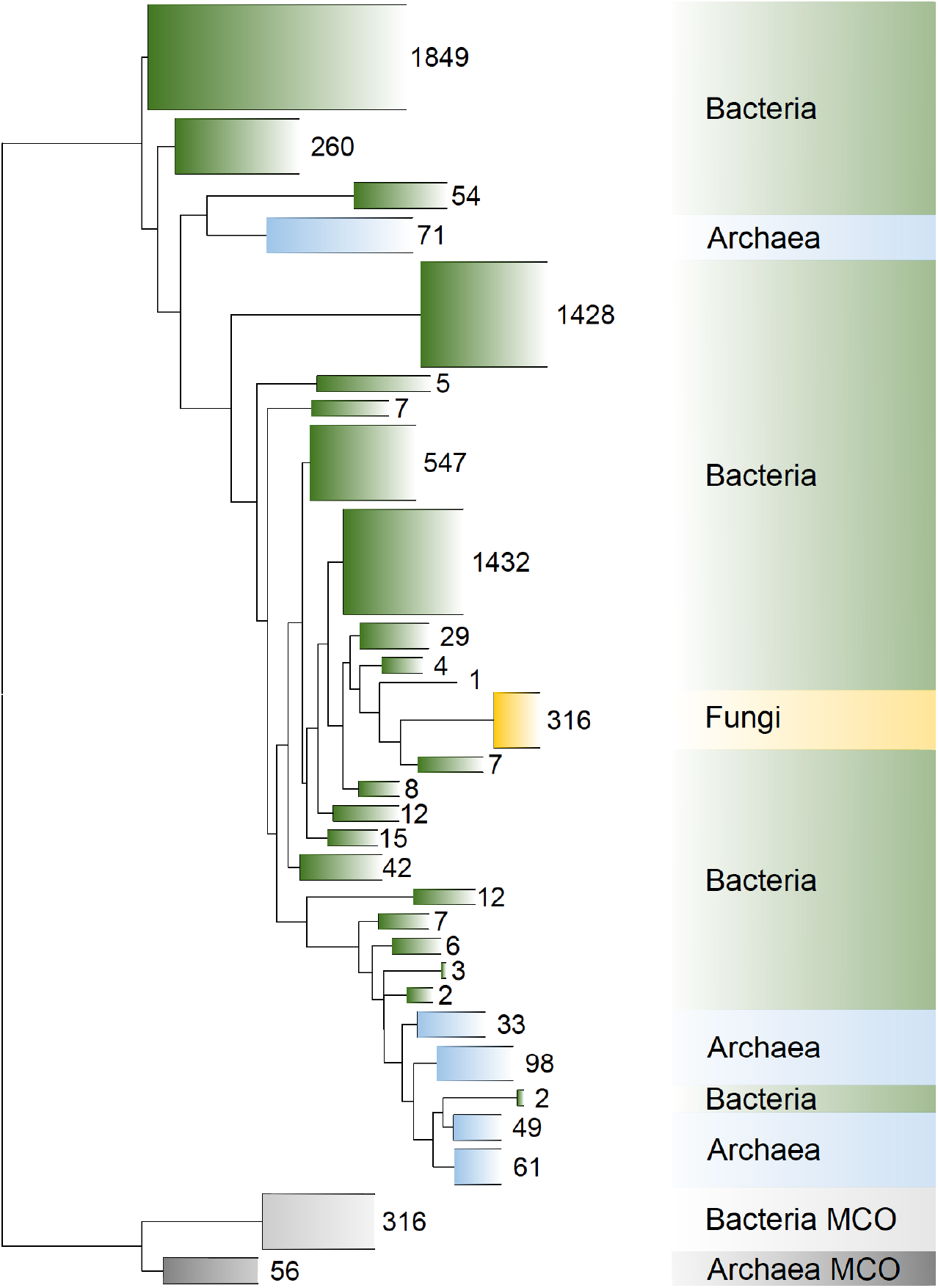
Reference phylogeny of prokaryotic and fungal *nirK* created from 6 732 full-length genomic *nirK* gene sequences. The phylogeny was determined based on maximum likelihood analysis of amino acid sequences of *nirK* using the LG+G substitution model, and subsequently used as the reference tree for phylogenetic placement of *nirK* gene fragments retrieved from metagenomes. Clades were collapsed with the number of unique species per collapsed clade indicated. The outgroups consist of multi-copper oxidase (MCO) sequences originating from cyanobacterial and thaumarcheotal genomes.

The reference phylogeny for the fungal 18S rRNA gene was obtained using the reference sequences and tree from SILVA (SSU Ref NR 99 138.1; [37]). The tree was first manually pruned in ARB to retain only representative species of each of the major eukaryotic lineages. The RNA sequences were then transformed to DNA prior to deduplication using BBMap (v.38.90) [38], resulting in 3 559 sequences in the tree.

### Screening metagenomes for *nirK* and 18S rRNA gene fragments

Fragments of *nirK* and 18S rRNA gene sequences were identified in metagenomes using GraftM (v. 0.13.1) [39]. GraftM utilizes custom gene reference packages to search metagenomes using HMMER followed by phylogenetic placement of the identified *nirK* gene fragments into a reference tree. For each gene, tree model parameters required for running GraftM were calculated using RaxML (v.1.1.0) [40] and the tree was re-rooted using FigTree (v.1.4.4) [41]. We used only forward reads and ran GraftM with default parameters except for restricting the read length to 150, resulting in screening only the first 150 nt of each read regardless of the total read length. Our *nirK* identification method was validated by generating a mock dataset of 24 670 150 nt fragments of *nirK* sequences present in the phylogeny, followed by GraftM searches using the specified parameters. Sensitivity was calculated by determining the fraction of eukaryotic *nirK* fragment sequences placed outside the fungal clade on the tree (13 117 of 13 266 reads, 98.8%). Specificity was determined using 1 427 775 fragments derived from 19 037 multi-copper oxidase family sequences which were identified in a database of bacterial and archaeal MAGs [42]. Only 0.05% (102 of 195 039) sequences placed in the tree were inappropriately placed within the fungal group.

### Phylogenetic and statistical analyses

We used a combination of the phylogenetic placement visualization and analysis tools guppy (v.1.1.alpha19-0-g807f6f3) [43] and gappa (v.0.8.0) [44] to remove non-target *nirK* and 18S rRNA gene fragments and classify *nirK* as being of bacterial, archaeal and fungal origin. Since GraftM classifies reads based on their placement in the phylogenetic tree, we kept the most likely placement of each *nirK* fragment (i.e. point mass) and counted it as belonging to the target microbial group, if the mass of possible placements for a fragment sequence reached a threshold of one for the selected clade The probability distribution of the most likely placements of each biome were visualized using the gappa examine ‘lwr’ function and indicated the probability of the most likely placement, given as a likelihood weight ratio (Fig. S1). Placements within trees were visualized using iTOL v5 [45] and to compare aggregation of placements within fungal clades or for species among biomes, the placements were visualized for each biome individually in iTOL. The trees were then visually inspected for clades or leaves with placements that were mostly restricted to a single biome and with multiple placements.

Because of the inherent problem with undersampling of functional genes in metagenomes having only few or zero gene fragment counts per metagenome [46], we excluded zero-valued samples and retained the 1 485 metagenomes that included both fungal *nirK* and fungal 18S rRNA gene fragment counts when reporting counts and comparing gene ratios across biomes. This minimizes zero-inflation within the dataset and subsequent skewing of the data, but bears the risk of overrepresentation of metagenomes with (multiple) gene fragment counts, resulting in inflated mean counts of functional genes.

All statistical analyses were carried out in the R environment (v.4.1.2). To account for differences in sequencing depth, fungal *nirK* abundance was normalized by dividing the number of *nirK* fragment counts by the total number of reads in the corresponding metagenome. The fungal *nirK* fragment counts were also divided by the number of fungal 18S rRNA gene sequences detected to account for differences in total fungal abundance across samples. To compare the fungal *nirK* abundance with its prokaryotic counterparts, the fungal/prokaryotic *nirK*, ratio was calculated. The effect of biomes on gene abundances was evaluated after negative log-normal transformation of the data. A generalized linear model approach was chosen utilizing a gamma distribution combined with a log link function (f*nirK*, f*nirK*/p*nirK*) or a Gaussian distribution (f*nirK*/18S) followed by analysis of variance (ANOVA), using the ‘stats’ (v.4.0.5) package. The Levene’s test was used to test for homogeneity of variance using the ‘car’ package [47] and the model was visually inspected and results reported in Table S2. Pairwise and multiple comparisons of biomes were carried out using the Dunn–Sidák correction method within the ‘emmeans’ package [48].

For the analysis of potential factors driving abundances of fungal *nirK*, ratios with overall fungal abundance and with prokaryotic *nirK*, a minimum of 25 metagenomes per biome were used to preserve statistical power. Further, the metadata associated with metagenomes containing fungal *nirK* was reduced to soil factors known to be relevant for denitrification (Table S3). These include different variables related to soil C and N content, soil texture and moisture known to affect oxygen levels, soil pH, and copper content, as NirK is a copper-dependent nitrite reductase [49, 50]. Spearman’s correlations were used to test for relationships between soil physicochemical variables and gene abundances or ratios using the packages ‘corrr’ [51] and ‘Hmisc’ [52]. Rhizosphere samples were excluded from this analysis since metadata was largely missing and fungal *nirK* abundance was instead compared across 14 host plant taxa.

## RESULTS

### Biome-related patterns of fungal denitrifier abundance

To investigate the prevalence of fungal denitrifiers in terrestrial ecosystems, metagenomes derived from terrestrial biomes representing 608 globally distributed sampling sites were analyzed (Fig. 1a). Based on phylogenetic placement of fragments of the fungal and prokaryotic denitrifier marker gene *nirK* in the metagenomes into a *nirK* reference tree (Fig. 2), archaeal, bacterial, and fungal *nirK* fragments were detected in 97, 100, and 76% of the metagenomes, respectively. Fungal *nirK* accounted for 4.5 ± 6.9 (mean ± SD) gene fragment counts per metagenome, when assigning the value “0” when fungal *nirK* was not detected. Fungal *nirK* counts were nearly 10 and 200 times lower than that for archaea and bacteria, respectively. By contrast, fungal 18S rRNA gene fragments were identified in 99% of the metagenomes, with an average of 457 ± 1097 reads per metagenome. The detection of fungal *nirK* differed among biomes, with the lowest proportion of metagenomes with zero fungal *nirK* fragments found in cropland metagenomes (2%) and the highest in deserts (37%) (Fig. S2a). These zero-counts may not indicate absence of fungal *nirK* but could potentially be the result of undersampling, as suggested by the lower number of total reads consistently recorded for biome-specific metagenomes in which fungal *nirK* was not detected (Fig. S2b). Hence, for the subsequent analyses we retained the 1 485 metagenomes containing both fungal *nirK* and fungal 18S rRNA gene fragment counts, which corresponds to 825 forest, 150 grassland, 140 cropland, 72 desert, 63 tundra, and 235 rhizosphere metagenomes. In this subset, fungal *nirK* accounted for 6.0 ± 7.4 (mean ± SD) gene fragment counts per metagenome.

When metagenomes were grouped according to biome classifications at Level 2 (Table S1), the total abundance of fungal *nirK*, normalized to the total number of reads to account for variation in sequencing depth, was highest in forests and lowest in tundra and rhizosphere (Fig. 1b). Grouping at a refined biome classification (Level 3) showed that the high fungal *nirK* abundance in forests was driven by the abundance in tropical and subtropical moist broadleaf forests (Table 1). The fungal *nirK* abundance was significantly higher in these biomes compared with Mediterranean forests, woodlands and shrublands, which were similar to deserts, tundra and rhizosphere. By contrast, the proportion of fungi carrying *nirK*, defined as the ratio of fungal *nirK* counts to those of the fungal 18S rRNA gene, was lower in forests and rhizosphere compared to other biomes (Fig. 1c). Comparisons at biome Level 3 revealed that the gene ratios were highest in tropical and subtropical dry broadleaf forests and lowest in Mediterranean forests and shrublands (Table 1).

**Table 1.**
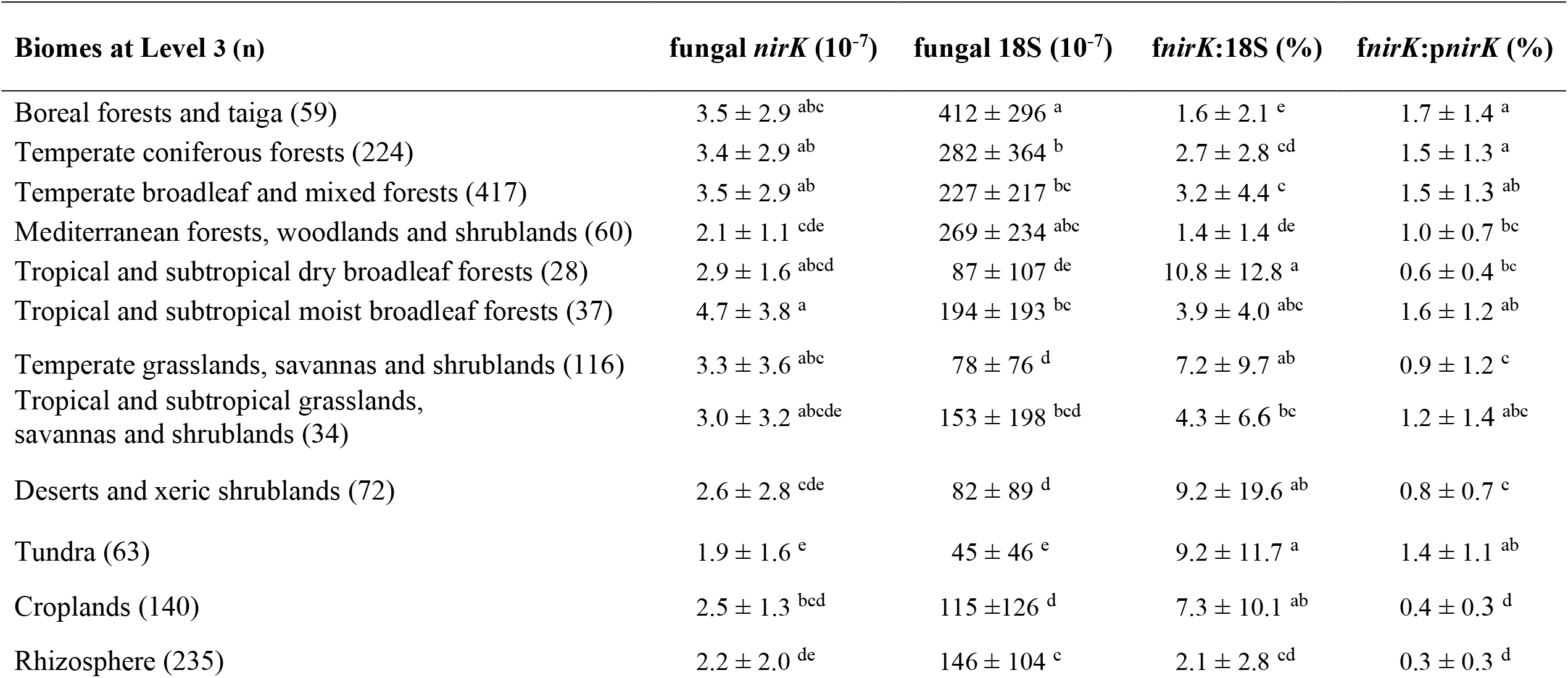
Abundance of fungal *nirK* (f*nirK*) and fungal 18S rRNA genes (18S) in terrestrial biomes at Level 3.

### Identity of fungal denitrifiers across biomes

The fungal *nirK* fragments collected from all terrestrial biomes spanned the fungal *nirK* reference tree, which is based on *nirK* from the classes Eurotiomycetes, Dothideomycetes, Leotiomycetes, Saccharomycetes, and Sordariomycetes (Fig. 3). The two largest classes, Eurotiomycetes and Sordariomycetes, were dominated by reference *nirK* sequences similar to those in the genera *Aspergillus, Fusarium*, and *Penicillium*. With two exceptions, *Fusarium nirK* sequences formed a monophyletic clade, whereas *Aspergillus nirK* sequences were split into several clades across the tree, interleaved by clusters of Eurotiomycetes members such as *Trichophyton, Paracoccidioides*, and *Blastomyces*. A large fraction of the metagenome *nirK* sequences was best placed in the region of the tree corresponding to *nirK* from species within the Eurotiomycetes. These include regions of the reference tree corresponding to *Aspergillus westerdijkiae* (2% of placements), *Chrysosporium tropicum* (1.7%), *Paracoccidioides brasiliensis* (1.4%), and several species of the genus *Exophiala* (1.6%). In addition, placements aligning to the single representative of the class Dothideomycetes, *Acidomyces richmondensis* (2.4%), were abundant. We also noted high *nirK* counts corresponding to *Antarctomycetes* (1.7%) and *Pseudogymnoascus* (4.9%) within the class Leotiomycetes. Among the Sordariomycetes, *nirK* fragments detected in all terrestrial biomes were frequently placed close to species of *Fusarium* (particularly *F. neocosmosporiellum* (3.4%)), *Dactylonectria* (2.5%), and *Scedosporium* (2.5%), as well as *Trichoderma hamatum* (0.9%), *Raffaelea lauricola* (4.5%), and *Purpureocillium lilacinum* (8.1%). A fraction of *nirK* sequences (18.6%) were placed at the basal part of the fungal tree.

**Fig. 3.**
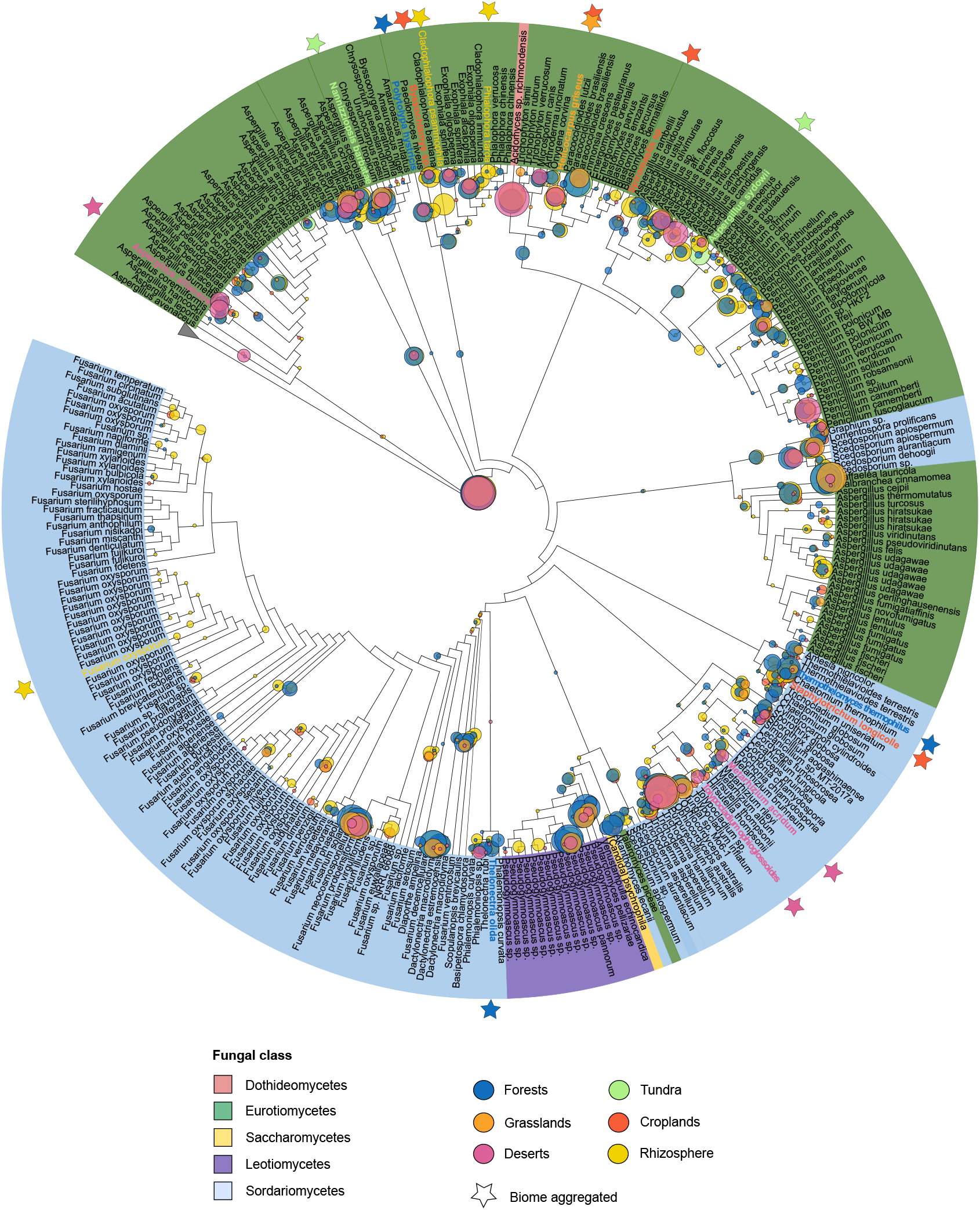
Phylogenetic placements of fungal *nirK* gene fragments detected in soil and rhizosphere biomes within the *nirK* reference cladogram tree. Leaf color indicates the fungal class, and the outgroup sequences are collapsed. The most likely phylogenetic placement for each read is represented by a circle and colored according to the biome classification at Level 2. The circle size indicates the number of placements on a given tree edge. Stars mark branches enriched in fungal *nirK* placements within a biome compared to other biomes. Biome-specific placements at Level 2 are shown in Figure S3.

Visual comparison of placements across biomes grouped at Level 2 showed forest metagenomes having more pronounced aggregations of placements within the genera *Polytolypa, Thermothelomyces*, and *Thelonectria* summing up to 0.7%, 1.5% and 0.6% of placements within forest biomes respectively (Fig. 3; Fig. S3). Placements of *Helicocarpus* (1.9%) were enriched in grasslands, whereas desert metagenomes were enriched in *nirK* from the mold *Aspergillus alliaceus* (2.2%), the genera *Tolypocladium* (1.1 %), and *Metarhizium* (1.1%). In tundra metagenomes, placements of *Nannizziopsis* (2.8%), and *Aspergillus sydowii* (3.3%) were found in comparably high abundance and in croplands, placements within the genera *Byssochlamys* (0.6%), *Helicocarpus* (1.4%), *Spiromastix* (0.4%), and *Staphylotrichum* (2.2%) were more prominent. In rhizospheres, the genera *Cladophialophora* (2.1%) and *Phialophora* (0.5%) were relatively abundant when compared to other biomes. Furthermore, placements assigned to *Fusarium oxysporium* were almost exclusively found in rhizosphere metagenomes (1%), whereas placements in this part of the tree were rare for fungal *nirK* detected in other biomes (Fig. 3; Fig S3).

### Environmental drivers of fungal *nirK* abundances

To determine environmental factors driving the prevalence of fungal denitrifiers, we related fungal *nirK* counts with metagenome associated edaphic variables. Across all terrestrial biomes combined, fungal *nirK* abundance was positively correlated with soil organic carbon (SOC), ammonium content, soil moisture, and clay content, but negatively associated with the carbon to nitrogen ratio (C/N), and pH (Fig. 4). The overall associations with C/N and pH were driven by the significant correlations with fungal *nirK* in forest biomes, and those with SOC by the significant correlations with fungal *nirK* in croplands. Across biomes classified at Level 2, non-significant or contrasting relationships were observed for some soil variables, but soil ammonium content, soil moisture, and clay content correlated positively with fungal *nirK* across several biomes. For soil N content, biome-specific relationships with fungal *nirK* were detected with positive correlations in croplands and negative in forests.

**Fig. 4.**
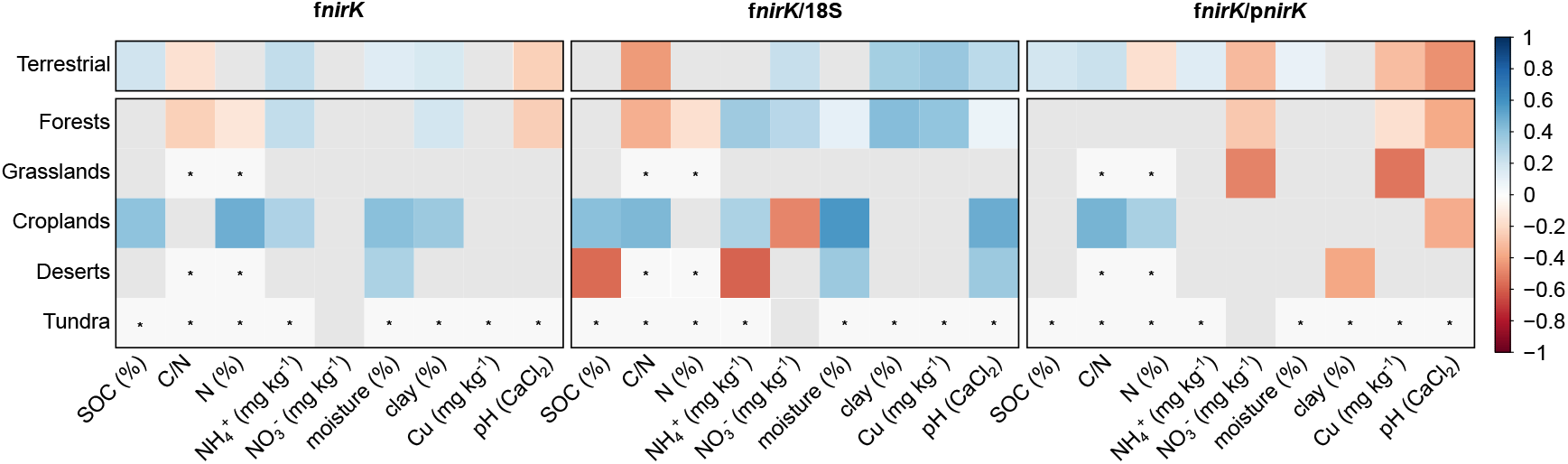
Spearman ranked correlations of fungal *nirK* abundance, and ratio to 18S rRNA gene abundance (f*nirK*:18S), and prokaryotic *nirK* (f*nirK*: p*nirK*) with edaphic variables. Correlations are shown for all terrestrial biomes and five terrestrial biomes at Level 2. Rhizosphere metagenomes were not included due to the absence of associated metadata. The correlations are colored if significant (p<0.05) according to their strength, with red for negative, blue for positive correlations. Non-significant correlations are colored in grey and those not determined due to insufficient sample number (n<25) are marked with an asterisk. Sample size differed among metagenomes as indicated in Table S3. SOC: soil organic carbon, C/N: total carbon to total nitrogen ratio, N: total nitrogen, NH_4_^+^ : ammonium, NO_3_^-^: nitrate, moisture: soil moisture, clay: soil clay content, Cu: soil copper content, pH: soil pH measured in CaCl_2_. NO_3_^-^ in tundra soils was reported as µM.

The fraction of the fungal community carrying *nirK* across all biomes was largely related to the same soil factors as the fungal *nirK* abundance, apart from pH which showed a positive relationship with the proportion of fungal *nirK* (Fig. 4). The relationship with pH was consistent for cropland, desert, and to a lesser extent, in forest soils. We also noted that the fraction of fungal *nirK* increased with increasing copper content, which was mainly driven by the forest soils. Among biomes, we detected contrasting associations with the proportion of fungal *nirK* in the fungal community and SOC, C/N, ammonium, and nitrate (NO_3_^-^). The overall decrease of the proportion of fungal *nirK* in relation to the C/N ratio was observed also in forests, whereas croplands showed a positive relationship. Ammonium was significantly correlated with the proportion of fungal *nirK* in both forest and cropland biomes, yet negatively correlated in the desert soils. Opposing patterns were also observed for NO_3_^-^ in forests and croplands, with negative associations in the latter. For rhizosphere metagenomes, for which only metadata on the host plant species was available, we noted significantly lower proportion of fungal *nirK* within the fungal community in the rhizosphere of the Poaceae species compared to the average of all rhizosphere metagenomes (Fig. S4a).

### Fungal to prokaryotic *nirK* ratio

A dominance of prokaryotic *nirK*, i.e. the sum of bacterial and archaeal *nirK* fragment sequence counts, was observed across all biomes (Fig. 1d). The highest ratio between fungal and prokaryotic *nirK* was found in forest and tundra soils, intermediate ratios were detected in grasslands and deserts, and the lowest in croplands and rhizosphere. Within forests, this ratio was significantly higher in boreal forests and taiga and temperate coniferous forests (Table 1). For the rhizosphere, the ratio between fungal and prokaryotic *nirK* was significantly lower in the metagenomes from *Miscanthus* sp. and *Populus* sp. when compared to the average of all rhizosphere samples (Fig. S4b).

## DISCUSSION

Here, we evaluated the distribution and abundance of fungal *nirK* in 1 980 metagenomes, providing a global survey of fungal denitrifiers in terrestrial ecosystems. Although fungal denitrifiers were overall rare compared to their archaeal and bacterial counterparts, they showed biome-specific differences in both abundance and species distributions. Our study further highlights that the dominant soil fungal denitrifiers are globally distributed organisms adapted to a broad range of climatic and environmental conditions. Known cosmopolitan fungal species carrying *nirK*, such as *Aspergillus westerdijkiae, A. sydowii, Penicillium solitum*, or *Fusarium neocosmosporiellum*, were identified in all the Level 2 biomes. Similarly, Dothideomycetes *nirK* sequences were detected across biomes, despite being represented by a single species (*Acidomyces richmondensis*) in the *nirK* reference phylogeny. Most members of the Dothideomycetes exhibit a saprotrophic lifestyle, but some are known as pathogens and endophytes [53]. *Aspergillus* and *Penicillium* species have been isolated from a range of environments [54, 55], and are known for their efficient dispersal strategies [56] and stress tolerance [57, 58]. Members of both genera also have the capacity to produce powerful extracellular enzymes for lignin and xylose degradation [59], an important trait supporting growth in environments with poor availability of easily accessible C substrates. In contrast to *Aspergillus*, the distribution of certain *nirK*-carrying *Fusarium* appears to be more biome specific, as members of the *Fusarium oxysporum* species complex were proportionally more abundant and diverse in the rhizosphere than in the other biomes. Increased fungal contribution to N_2_O production by denitrification has been assigned to rhizosphere processes [60] and *F. oxysporum* in particular is known to produce substantial amounts of N_2_O [5, 61]. *F. oxysporum* is also a typical root pathogen, suggesting that host-pathogen interactions might play a role for increased fungal denitrifier abundance. Accordingly, there are indications that the fungal nitric oxide reductase P450nor is also involved in fungal virulence [62]. Further evidence for the relevance of host-pathogen interactions as a driver of fungal denitrifiers is the relatively high proportion of pathogenic fungi in the reference phylogeny, and our observation that some of these pathogenic fungi were more associated with certain biomes, based on the placements of the metagenome sequences. This includes the genera *Tolypocladium* and *Metarhizium*, both known entomopathogens in warm deserts, the plant pathogen species *Byssochlamys*, and *Thelonectria* which infests hardwood in forests. Overall, this suggest that plant-pathogen interactions support fungal denitrifiers, and a possible mechanism is that denitrification increases virulence, as shown for several bacteria. [63].

The majority of the placements were located close to the leaves of the reference tree, suggesting that the majority of the metagenome-derived fungal denitrifiers are similar to known fungal denitrifiers. Despite uncertainty in the placement of fungal *nirK* reads on the reference phylogeny, we detected many of the dominant fungal taxa carrying *nirK* that were observed previously [64–66]. Nevertheless, PCR-based approaches specifically targeting *nirK* reveal a more diverse community of soil fungal denitrifiers [67] than we observed in the metagenomes, where the presence of fungal *nirK* was rare and the majority being similar to known fungal denitrifiers. Notably, *nirK* sequences obtained by amplicon sequencing of arable soil sampled from a long-term field experiment showed more basal placements of fungal *nirK* in the phylogeny than those detected in the metagenomes [67], underlining that the metagenomic *nirK* sequences mainly capture the most dominant of fungal denitrifiers due to limited sequencing depth. Still, 18.6% of the placements were found at the most basal part of the fungal tree, which implies the presence of eukaryotic organisms with *nirK* not captured by our reference phylogeny. These placements could point to unknown fungal denitrifiers, non-fungal eukaryotes such as some algae [21] which carry *nirK* but are not represented in the phylogeny, or sequences from highly conserved portions of the alignment and which are therefore difficult to parse. Nonetheless, the metagenomes captured the dominant terrestrial fungal denitrifiers, contributing to a better understanding of their diversity, distribution and ecology across global soils.

The abundance of fungal *nirK* in specific biomes was in general not explained by the overall abundance of fungi. Correlations with soil factors indicate that fungal denitrifiers are most abundant under conditions generally favorable for denitrification [4], with high moisture, and C and N availability, although their abundance increased with decreasing pH. Low pH has been shown to stimulate fungal denitrification [19] and Xu et al. [65] found a significant pH effect on fungal *nirK* abundance. Nevertheless, our results indicate that fungal denitrifiers appear to thrive at higher pH compared to fungi in general and may also be less tolerant to dry conditions than the overall fungal community. Alternatively, increased moisture would indicate decreased soil aeration, which could promote fungal denitrifiers due to their capacity for facultative respiration under anoxic conditions [68]. Other relationships between the proportion of fungal *nirK* in the fungal community and soil properties displayed biome-specific patterns, likely because variation of these properties differed across biomes. For example, increasing inorganic N content generally promoted denitrifying fungi relative to the overall fungal community, but negative associations to ammonium and nitrate were found in deserts and croplands, respectively. Because of losses by volatilization, desert soils often have lower ammonium levels than forest and cropland soils [69]. Increased ammonium levels in deprived desert soils have resulted in depression of the fungal order Sordariales [70], and indeed, a large fraction of *nirK* sequence fragments found in desert soil metagenomes were classified as Sordariales, supporting a negative relationship between the proportion of fungal denitrifiers within the fungal community and ammonium. The negative association with soil nitrate and the proportion of denitrifying fungi in croplands, despite the positive correlation in forests, aligns with the higher nitrate levels in croplands. This could indicate a nitrate threshold causing restructuring of the fungal community. Similarly, plants might affect the proportion of fungal denitrifiers in the rhizosphere, as observed for the lower abundance of fungal *nirK* with Poaceae, but due to lack of data on the environmental conditions in the rhizosphere, it cannot be concluded whether these effects are host or soil driven.

Fungal denitrifiers relative to their prokaryotic counterparts could be more important in tundra and forest soils, in particular boreal and temperate coniferous forests, compared to the other biomes. The lower pH in these systems, the negative correlation of the ratio between fungal and prokaryotic *nirK* fragment counts with pH, and previous reports of fungal denitrification prevailing in acidic soils [18, 19] indicate pH as a relevant predictor. In croplands, increasing C/N is another possible driver, supported by Chen et al. [15], who reported that complex organic C substrates enhanced fungal denitrification compared to that of bacteria. Nevertheless, even under the most favorable conditions, the abundance of fungal denitrifiers relative to the prokaryotes remains low and constitutes approximately 1% of all *nirK*-type denitrifiers. We based our counts of prokaryotic denitrifiers on *nirK* alone, as the other nitrate reductase among denitrifiers, NirS, is much less abundant than NirK in terrestrial ecosystems [71–74]. Hence, adding *nirS* counts would have limited impact on the ratio between fungal *nirK* and prokaryotic *nir* genes. Nonetheless, the low relative abundance of fungal *nirK* detected across the metagenomes, indicates their potential contribution to denitrification and previously reported abundance based on PCR-based approaches may have been overestimated. Fungal *nirK* quantification by quantitative PCR without sequence correction has recently been shown to be unreliable as it overestimates the abundance of fungal denitrifiers by orders of magnitude [66, 67]. Similarly, there is evidence that selective inhibition approaches commonly used to discriminate between fungal and prokaryotic denitrification, overestimate fungal contribution to denitrification [75]. Overall, their role in denitrification and contribution to soil N_2_O emissions may therefore be less important than previously suggested.

Despite having a minor role in global denitrification, it remains intriguing why some fungi have the ability to denitrify. One explanation could be related to the saprotrophic and opportunistic pathogenic lifestyles that known denitrifying fungal taxa exhibit, which also involves nitric oxide detoxification [76] resulting from NO-biosynthesis under nitrosative stress caused by the host response during fungal infection [77]. Denitrification may also increase fungal virulence [62, 78] as discussed previously. For saprotrophs and opportunistic pathogens, metabolic flexibility and stress tolerance are advantageous, and denitrification further allows them to stay metabolically active in oxygen-depleted environments, such as host tissue. Our finding that many fungal denitrifiers are stress-tolerant cosmopolitans makes fungal denitrifiers potential beneficiaries of global climate change [79], with a potential positive feedback through emissions of N_2_O. Moreover, warmer temperatures have been shown to increase soil-borne fungal plant pathogens and projections under different warming scenarios suggest an increase of these pathogens worldwide [80]. Future work should aim at gaining a deeper understanding of the importance of this understudied fraction of the denitrifying microbial community in a global change context.

## ACKNOWLEDGEMENTS

This work was supported by The Swedish Research Council Formas (grant number 2016-00477 to SH) and a career grant from the Swedish University of Agricultural Sciences to SH. Running BUSCO was enabled by resources provided by the Swedish National Infrastructure for Computing (SNIC) at UPPMAX partially funded by the Swedish Research Council through grant agreement no. 2018-05973. We would like to acknowledge the contribution of the Biomes of Australian Soil Environments (BASE) consortium in the generation of data used in this publication. The BASE project was supported by funding from Bioplatforms Australia through the Australian Government National Collaborative Research Infrastructure Strategy (NCRIS).

## AUTHOR CONTRIBUTIONS

S.H. and C.M.J. designed the study. C.M.J. and A.S. created the database with support from G.P. and Y.B. C.M.J., G.P. and Y.B. created the *nirK* phylogeny. Y.B. and G.P. analyzed the data. Y.B., G.P. and S.H. wrote the paper with contributions from all authors. All authors read and approved the final version of the manuscript.

