## Supplemental information for "Distribution and environmental drivers of fungal denitrifiers in global soils"

**Table S1. Biome classification, number of metagenomes per biome and references for metagenomes included in the study.** The number (n) of metagenomes per reference is indicated. The total number of metagenomes per biome Level 2 and 3 are shown in parentheses. Unpublished metagenomes are indexed with their NCBI BioProject number (PRJNA). Full references are provided in the supplementary "Reference" section.

| Biome Level 2 | Biome Level 3 | n | Reference |
|---|---|---|---|
| Croplands (144) | Croplands (144) | 41 | Bissett et al. 2016 [1] |
| | | 32 | Hartman et al. ISME J 2017 [2] |
| | | 3 | Mendes et al. ISMEJ 2018 [3] |
| | | 12 | Orellana et al. Appl Env Mic 2018 [4] |
| | | 36 | PRJNA717057 |
| | | 20 | Xu et al. Nat Com 2020 [5] |
| Deserts (115) | Deserts and Xeric Shrublands (115) | 3 | Bahram et al. Nature 2018 [6] |
| | | 39 | Bissett et al. 2016 [1] |
| | | 73 | NEON, 2021 [7] |
| Forests (1118) | Boreal Forests & Taiga (89) | 7 | Bahram et al. Nature 2018 [6] |
| | | 61 | NEON, 2021 [7] |
| | | 21 | Wilhem et al. Sci Data 2017 [8] |
| | Mediterranean Forests Woodlands and Scrub (84) | 17 | Bahram et al. Nature 2018 [6] |
| | | 45 | Bissett et al. 2016 [1] |
| | | 22 | NEON, 2021 [7] |
| | Temperate Broadleaf and Mixed Forests (542) | 66 | Bahram et al. Nature 2018 [6] |
| | | 139 | Bissett et al. 2016 [1] |
| | | 292 | NEON, 2021 [7] |
| | | 12 | Sorensen et al. Nat Microbiol 2019 [9] |
| | | 21 | Wilhem et al. Sci Data 2017 [8] |
| | | 12 | Xiao-Jun Allen Liu; unpublished, PRJNA621569, PRJNA621570, PRJNA654925 –34 |
| | Temperate Conifer Forests (273) | 8 | Bahram et al. Nature 2018 [6] |
| | | 60 | Diamond et al. Nat Microbiol 2019 [10] |
| | | 161 | NEON, 2021 [7] |
| | | 44 | Wilhem et al. Sci Data 2017 [8] |
| | Tropical and Subtropical Coniferous Forests (3) | 3 | Bahram et al. Nature 2018 [6] |
| | Tropical and Subtropical Dry Broadleaf Forests (41) | 41 | NEON, 2021 [7] |
| | | 78 | Bahram et al. Nature 2018 [6] |

| | | | |
|---|---|---|---|
| | | 6 | Bissett et al. 2016 [1] |
| | Tropical and Subtropical Moist Broadleaf Forests (86) | 2 | Mendes et al. ISMEJ 2017 [3] |
| Grasslands (241) | Montane Grasslands and Shrublands (5) | 2 | Bahram et al. Nature 2018 [6] |
| | | 3 | Bissett et al. 2016 [1] |
| | Temperate Grasslands Savannas and Shrublands (194) | 2 | Bahram et al. Nature 2018 [6] |
| | | 13 | Bissett et al. 2016 [1] |
| | | 179 | NEON, 2021 [7] |
| | Tropical and Subtropical Grasslands Savannas and Shrublands (42) | 13 | Bahram et al. Nature 2018 [6] |
| | | 29 | Bissett et al. 2016 [1] |
| Tundra (89) | Tundra (89) | 5 | Bahram et al. Nature 2018 [6] |
| | | 27 | NEON, 2021 [7] |
| | | 57 | Woodcroft et al. Nature 2018 [11] |
| Rhizosphere (273) | *Amaranthus* sp. | 13 | Bandla et al. Scientific Data 2020 [12] |
| | *Arabidopsis thaliana* | 49 | Levy et al. 2018 Nature Genetics [13] |
| | *Asparagus* sp. | 12 | Crovadore et al. MRA 2017 [14] |
| | *Phaseolus vulgaris* | 23 | Mendes et al. ISMEJ 2018 [3] |
| | *Brassica alboglabra* | 16 | Bandla et al. Scientific Data 2020 [12] |
| | *Brassica parachinensis* | 15 | Bandla et al. Scientific Data 2020 [12] |
| | *Citrus* sp. | 23 | Xu et al. Nature Communications 2018 [15] |
| | *Colobanthus quitensis* | 3 | Molina-Montenegro Polar Biology 2019 [16] |
| | *Colobanthus quitensis* + *Deschampsia antarctica* | 3 | Molina-Montenegro Polar Biology 2019 [16] |
| | *Zea mays* | 32 | PRJNA330341-47, PRJNA367156-68, PRJNA405457, PRJNA406023-27, PRJNA444376-80 |
| | *Gossypium* sp. | 1 | Singh et al. MRA 2020 [17] |
| | *Miscanthus* sp. | 43 | PRJNA330359-60, PRJNA365493-99, PRJNA366147-53, PRJNA366178-79, PRJNA367152 -53, PRJNA375575-80, PRJNA405458-61, PRJNA444381-85 |
| | *Populus* sp. | 13 | Blair et al. mSystems 2018 [18] |
| | *Helianthus annuus* | 1 | Babalola et al Data in Brief 2020 [19] |
| | *Panicum virgatum* | 25 | PRJNA330352-58, PRJNA365487-92, PRJNA375569-74, PRJNA405463-67, PRJNA444386 |
| | *Taxus cuspidata* | 1 | Hao et al. J. Basic Microbiology 2018 [20] |

**Table S2. Analysis of variances (ANOVA) of fungal *nirK* abundance in biomes at level 2.** The analysis was performed using a generalized linear model approach using a gamma distribution (f*nirK* and f*nirK*/p*nirK*) with a log link function and a gaussian distribution (f*nirK*/18S) after data transformation (negative log-normal). The F-distribution values for the Biome 2 response variable are indicated with the degrees of freedom in lower case letters.

| Abundance | Response | Deviance | F-value | P-value |
|---|---|---|---|---|
| f*nirK* | Biome 2 | 0.20 | $F_{5, 1479} = 17.74$ | $<2.2 \times 10^{-16}$ |
| f*nirK*/p*nirK* | Biome 2 | 16.19 | $F_{5, 1479} = 129.25$ | $<2.2 \times 10^{-16}$ |
| f*nirK*/18S | Biome 2 | 225.92 | $F_{5, 1479} = 35.26$ | $<2.2 \times 10^{-16}$ |

**Table S3. Soil variables used for correlation analysis with observed *nirK* abundance at biome level 2 (mean ± SD).** Comparison across biomes was performed using a generalized linear model approach with Sidak-pairwise comparisons. Different letters indicate significant differences across biomes. The number of metagenomes (n) associated with each biome and edaphic variable is indicated and the total number of metagenomes is indicated in parenthesis for each biome. SOC: soil organic carbon, C/N: total carbon to total nitrogen ratio, N: total nitrogen, $NH_4^+$: ammonium, $NO_3^-$: nitrate, moisture: soil moisture, clay: soil clay content, Cu: soil copper content, pH: soil pH measured in $CaCl_2$.

| **Biome** | **SOC (%)** | **n** | **C/N** | **n** | **Tot. N (%)** | **n** | **$NH_4^+$ (mg kg$^{-1}$)** | **n** | **$NO_3^-$ (mg kg$^{-1}$)** | **n** | **$NO_3^-$ (µM)** | **n** | **Moisture (%)** | **n** | **Clay (%)** | **n** | **Cu (mg kg$^{-1}$)** | **n** | **pH ($CaCl_2$)** | **n** |
|---|---|---|---|---|---|---|---|---|---|---|---|---|---|---|---|---|---|---|---|---|
| Forests (825) | 7.5 ± 10.7 $^{B}$ | 415 | 20.8 ± 7.2 $^{A}$ | 237 | 0.47 ± 0.47 $^{B}$ | 237 | 9.8 ± 13.5 $^{A}$ | 178 | 6.4 ± 12.0 $^{B}$ | 189 | 585.6 ± 2.3 $^{A}$ | 11 | 55.3 ± 113.3 $^{B}$ | 664 | 16.0 ± 11.8 $^{AB}$ | 173 | 1.0 ± 2.6 $^{B}$ | 178 | 4.9 ± 1.0 $^{C}$ | 680 |
| Grasslands (150) | 1.6 ± 1.3 $^{C}$ | 47 | 14.3 ± 2.0 $^{AB}$ | 9 | 0.24 ± 0.13 $^{AB}$ | 9 | 2.6 ± 1.9 $^{B}$ | 33 | 2.3 ± 1.8 $^{B}$ | 33 | NA | 0 | 16.2 ± 11.5 $^{C}$ | 131 | 14.0 ± 12.1 $^{AB}$ | 32 | 0.7 ± 0.6 $^{B}$ | 33 | 6.5 ± 1.0 $^{B}$ | 144 |
| Desert (72) | 1.3 ± 1.5 $^{C}$ | 37 | 17.6 ± NA | 1 | 0.43 ± NA | 1 | 4.9 ± 6.7 $^{B}$ | 36 | 4.9 ± 6.7 $^{AB}$ | 36 | NA | 0 | 4.6 ± 5.3 $^{C}$ | 61 | 12.6 ± 8.2 $^{B}$ | 36 | 0.7 ± 0.7 $^{B}$ | 36 | 7.0 ± 0.9 $^{A}$ | 71 |
| Tundra (63) | 25.3 ± 15.8$^{A}$ | 10 | 21.6 ± 3.7 $^{A}$ | 10 | 1.20 ± 0.79 $^{A}$ | 10 | NA | 0 | NA | 0 | 2.6 ± 2.3 $^{B}$ | 40 | 314.4 ± 173.7 $^{A}$ | 14 | NA | 0 | NA | 0 | 4.7 ± 0.4 $^{C}$ | 13 |
| Croplands (140) | 6.1 ± 7.5$^{B}$ | 93 | 13.9 ± 2.1 $^{B}$ | 32 | 0.74 ± 0.53 $^{AB}$ | 43 | 6.6 ± 7.4 $^{AB}$ | 84 | 9.1 ± 11.9 $^{A}$ | 84 | NA | 0 | 27.5 ± 21.0 $^{BC}$ | 83 | 21.2 ± 14.6 $^{A}$ | 48 | 2.6 ± 2.7 $^{A}$ | 41 | 7.1 ± 1.0 $^{A}$ | 41 |
| Rhizosphere (235) | NA | 0 | NA | 0 | NA | 0 | NA | 0 | NA | 0 | NA | 0 | NA | 0 | NA | 0 | NA | 0 | NA | 0 |

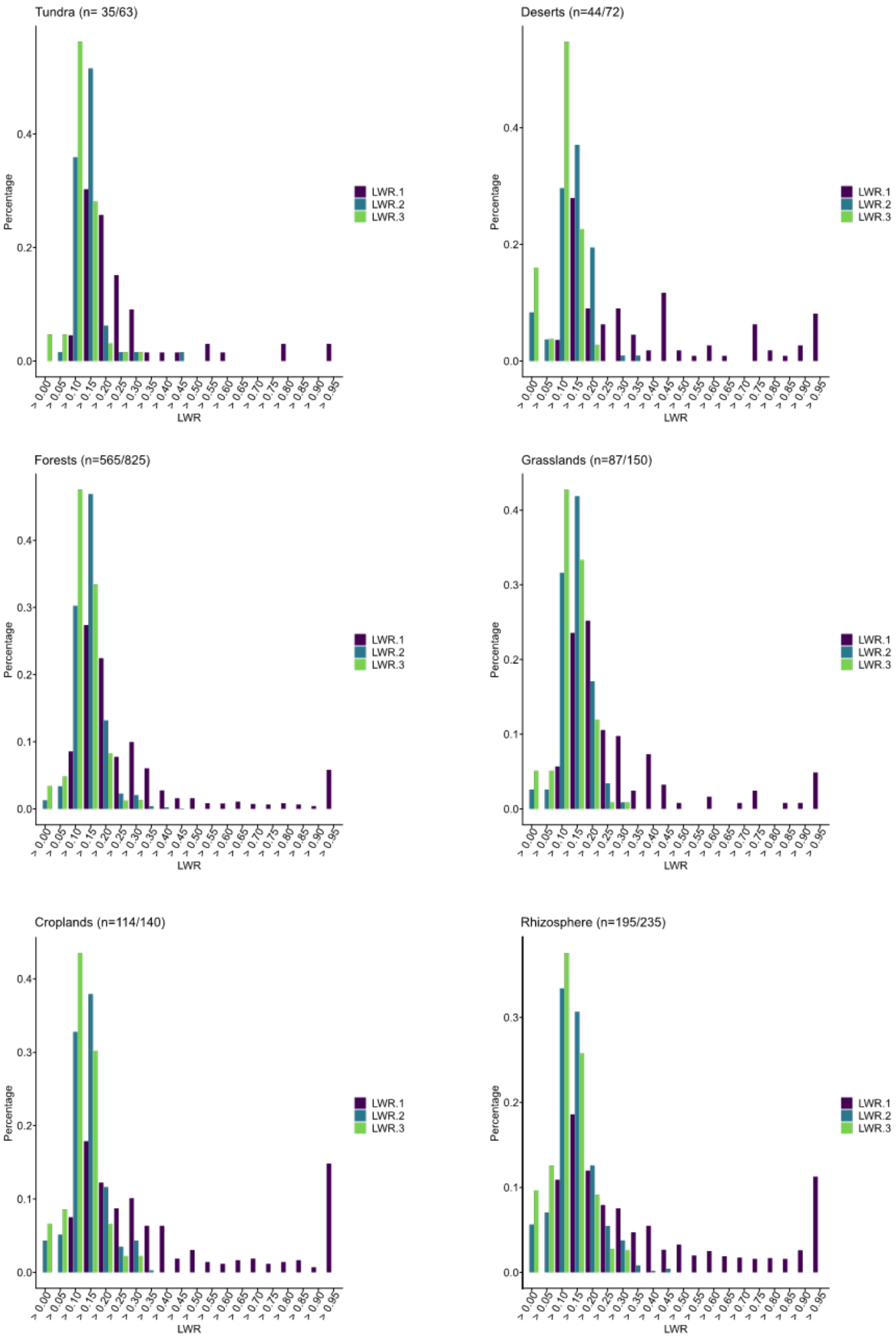


**Fig. S1. Histogram of the percentage of first, second and third most likely placements of *nirK* gene fragments from metagenomes at biome Level 2 in the *nirK* reference phylogeny.** The fractions of placements at each level as percentage is shown for different likelihood weight ratios (LWR), i.e. the certainty of a placement in the *nirK* reference phylogney. The different likelihood levels from most likely (LWR1) followed by the second (LWR2) and third most likely (LWR3) placement are shown next to each other. For analysis of abundance and correlations, only LWR1 placements were considered. Uncertainties of placements are expected due to for example sequencing errors and chimeric sequences, and missing reference sequences in the reference phylogeny, see Czech *etal*. 2022 [21].

**a**

| Biome 2 | MG's Total | MG's with fnirK | Zero - Counts (%) |
|---|---|---|---|
| Croplands | 144 | 141 | 2.1 |
| Deserts | 115 | 72 | 37.4 |
| Forests | 1118 | 834 | 25.4 |
| Grasslands | 241 | 156 | 35.3 |
| Tundra | 89 | 64 | 28.1 |
| Rhizosphere | 273 | 236 | 13.6 |
| Total | 1980 | 1503 | 24.1 |

**b**

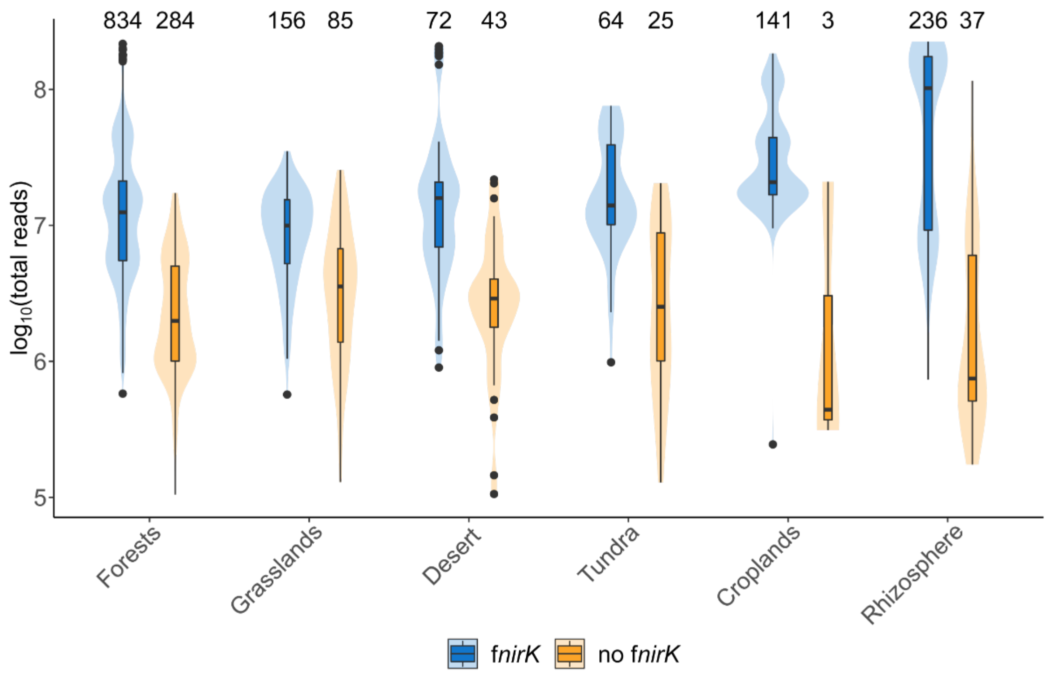


**Fig. S2. Distribution of metagenome size and number of metagenomes with zero fungal *nirK* fragement counts across biomes. a** Total number of metagenomes (MG's) per biome at Level 2 processed with GraftM and the total number of metagenomes with fungal *nirK* fragment counts included in the subsequent analyses . The fraction (%) of MGs without fungal *nirK* fragments detected is shown as zero-counts. **b** The total number of reads per metagenome ($log_{10}$ transformed) for each biome at Level 2 are split according to weather fungal *nirK* was detected or not. Box limits represent the inter-quartile range (IQR) with median values represented by the centreline. Whiskers represent values ≤1.5 times the upper and lower quartiles, while points indicate values outside this range. The shaded areas show kernel density estimations indicating the distribution of the data.

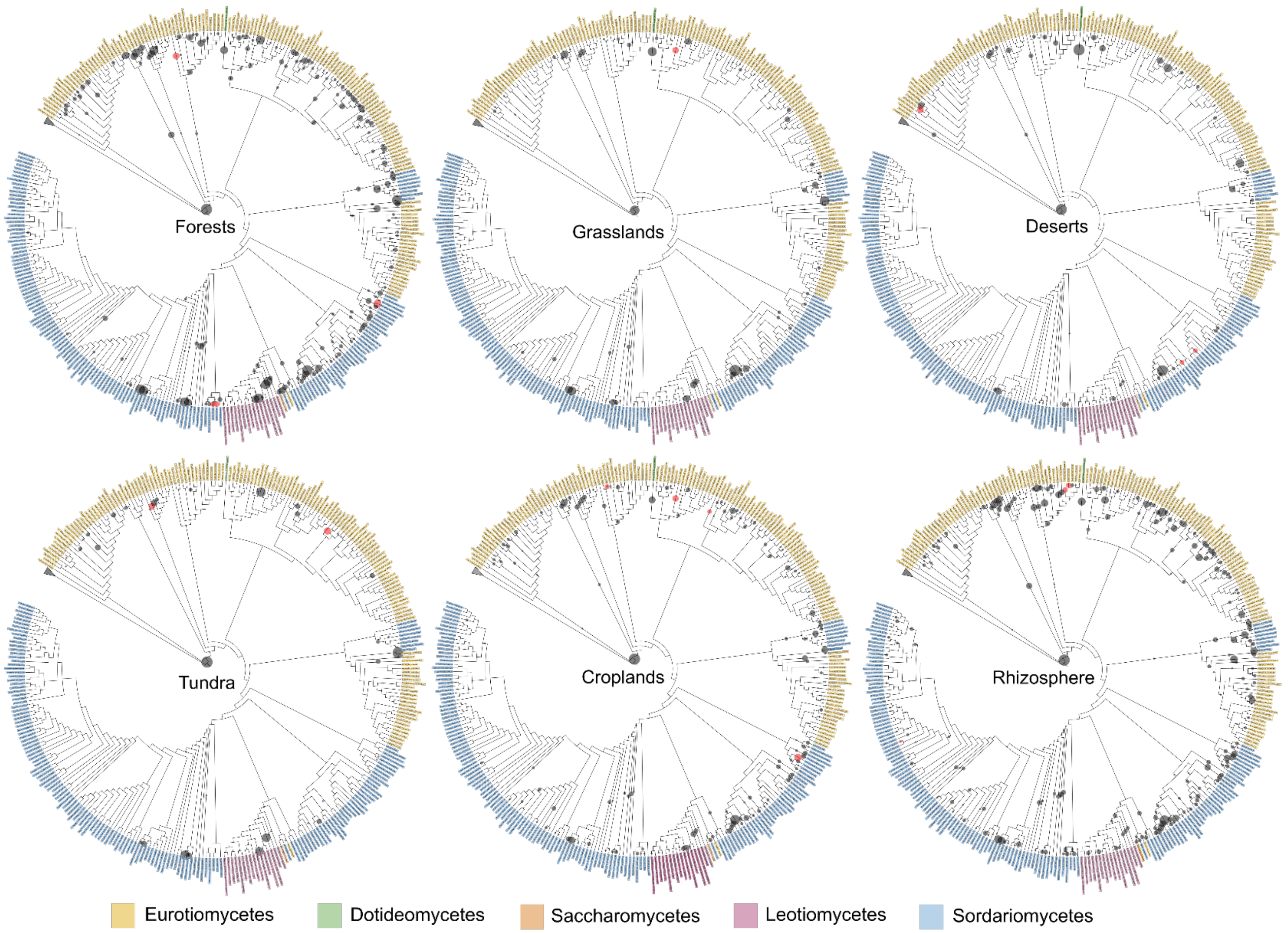


**Fig. S3. Phylogenetic placements of fungal *nirK* gene fragments within the *nirK* reference cladogram for each terrestrial biome at classified at Level 2**. Leaf color indicates the fungal class and the outgroup sequences are collapsed. The most likely phylogenetic placement for each read is represented by a circle and the size indicate the number of placements on a given branch. Red circles correspond to biome-aggregated fungal *nirK* placements, i.e. placements nearly exclusively found in one biome or with higher placement aggregations compared to other biomes.

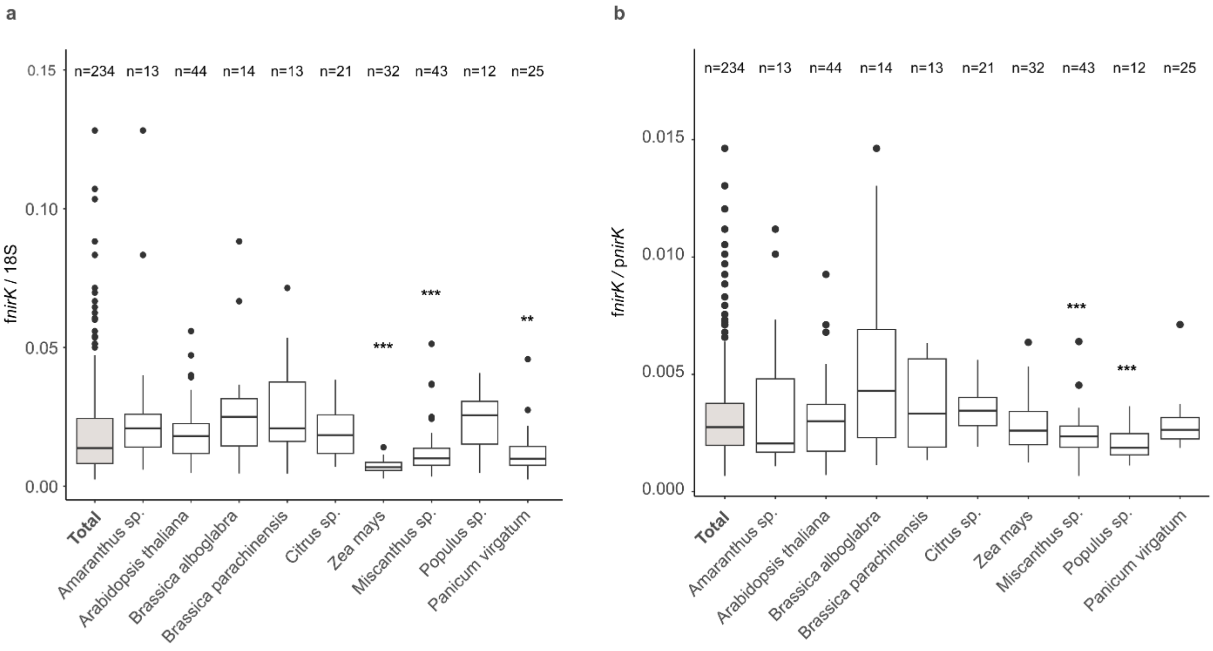


**Fig. S4. Comparison of fungal *nirK* counts in host species with n>10 metagenomes relative to the total of counts in rhizosphere. a** Counts of fungal *nirK* (f*nirK*) fragments relative to fungal 18S rRNA gene fragments (18S). **b** Counts of fungal *nirK* relative to prokaryotic *nirK* (p*nirK*). Stars represent significant (*$0.01<p<0.05$; **$0.001<p<0.01$; *** $p<0.001$) differences of the mean of each host species compared to the total of rhizosphere metagenomes determined by a two-sided t-test. Box limits represent the inter-quartile range (IQR) with median values represented by the centreline. Whiskers represent values ≤1.5 times the upper and lower quartiles, while points indicate values outside this range.
